# First insight into genetic diversity of two sympatric marten species between the Alps and Adriatic islands

**DOI:** 10.1101/2025.07.25.666739

**Authors:** Elena Buzan, Luka Duniš, Tilen Komel, Boštjan Pokorny, Carlos Rodríguez Fernandes, Zoran Marčić, Magda Sindičić

## Abstract

Closely related species occupying the same geographical area may exhibit markedly different genetic patterns due to differences in evolutionary history, ecology and behaviour. In this study, a landscape genetics approach is applied to investigate the genetic structure, diversity, and connectivity of two sympatric carnivore species, i.e. the European pine marten (*Martes martes*) and the stone marten (*Martes foina*) in Croatia and Slovenia. We analysed mitochondrial DNA sequences for both species and additionally used nuclear microsatellite markers for the pine marten. A total of 211 individuals (29 pine martens and 182 stone martens) from both mainland and island populations were analysed.

For pine marten, we found a significant genetic structuring, with pronounced differentiation between island and mainland populations, and a further substructure within the mainland. No significant isolation by distance was detected (Mantel test, *p* = 0.15), suggesting that genetic differentiation is driven more by habitat discontinuities and anthropogenic barriers rather than geographical distance alone. In contrast, stone marten exhibited weak genetic structure and high genetic diversity, indicating gene flow and potential landscape permeability for this more synanthropic species. These contrasting patterns underscore species-specific responses to landscape fragmentation and highlight the need to tailor management strategies accordingly.

## Introduction

Pine marten (*Martes martes*) and stone marten (*Martes foina*) are medium-sized mustelids that are very similar in terms of morphology, habitat preferences, foraging behaviour, and distribution [1–3]. Despite their wide range overlap and coexistence, encompassing most of central and southern Europe [4], creating favourable conditions for competition, few studies have analysed their spatial genetic distribution in sympatry [5]. Although stone marten is generally more adaptable in its habitat requirements and can shift to rural and urban areas, both species prefer forest habitats and tend to avoid open areas [6–8]. A recent study has shown that stone marten has a higher tolerance to human disturbance [9], suggesting that pine marten may dominate in the most suitable forest habitats. Although pine marten has long been considered a forest specialist, it has recently been repeatedly reported in rural areas [2,10,11], where it seems to exclude stone marten from residual forest fragments [9]. Stone marten is generally found in a variety of habitats, such as forested areas (deciduous forests, forest edges), steppes, rocky and cultivated areas. In south-western Europe, this species usually occurs in forests, while in central and north-eastern Europe it is more common in urbanised areas and mosaics of forest and field patches [3,7,12]. In contrast, the habitat of pine marten is generally associated with deciduous, mixed and coniferous forest, as well as scrubs [1,5,13].

So far, relatively few studies have addressed the genetics of pine and stone martens, mainly focusing on genetically identifying and distinguishing them in the field [5,14–19], with some looking into genetic diversity, population structure, and phylogeographic patterns, using nuclear markers [12,20,21], mitochondrial DNA [5,22–24], or a combination of both [25–28].

Both species exhibit complex phylogeographic structures that can be attributed to postglacial recolonization from refugia [24,29]. The Balkans served as a refugial area for the stone marten population, and in the absence of distinct geographical barriers, after the Last Glacial Maximum (LGM) martens expanded from the Balkan Peninsula to western Turkey [27], most of Europe, and subsequently into Central Asia [24,25] found that Bulgarian populations are part of a broader European lineage, suggesting historical gene flow and limited regional differentiation. In Greece, Tsoupas et al. identified high mitochondrial diversity and unique haplotypes, supporting the hypothesis that Greece may have acted as an additional glacial refugium [23]. In Turkey, İbiş et al. detected distinct genetic lineages within Anatolia, reinforcing the idea that the region harboured multiple refugia for stone marten during the LGM [22]. Building on this, Arslan et al. reported the existence of two refugia in Turkey, separated by the Taurus Mountains, corresponding to the western and eastern regions of the country [27]. They identified two distinct lineages: one originating from the eastern Anatolian refugium and the other from a western Anatolian or Balkan refugium. The western lineage colonized large parts of Europe after the LGM, including central and western Europe, Greece, and Bulgaria. In contrast, the eastern lineage appears to be endemic to Anatolia and may have reached the Iberian Peninsula through a human-mediated translocation.

Microsatellite studies of the two species have revealed significant insights into their genetic diversity and structure and dispersal patterns at smaller spatial scales. Wereszczuk et al. demonstrated that pine marten in Poland expanded from the south-western to the north-eastern regions in response to environmental changes, including climate warming [21]. In Portugal, Basto et al. identified northern, southern and south-eastern genetic groups within the country, with rivers acting as geographical barriers influencing their distribution [30]. Kyle et al. investigated the genetic diversity of the European pine marten using microsatellite markers across various locations, including England, Scotland, Ireland, Italy, Germany, Latvia, the Netherlands, Sweden, and Finland [31]. Their study revealed that continental populations exhibited higher genetic structure and lower genetic variation compared to their North American counterpart, i.e. American marten (*Martes americana*), which is attributed to habitat fragmentation and persecution. Island populations, such as those in Scotland and Ireland, showed significant genetic differentiation with lower genetic diversity. Interestingly, the English population displayed relatively high genetic diversity, possibly due to introgression with *M. americana*, indicating potential hybridization. Ruiz-González et al. conducted a comprehensive landscape genetic study of pine marten in the Basque Country in northern Spain, finding that gene flow between populations was influenced by natural vegetation and restricted by anthropogenic structures such as urban areas and roads [20]. Their findings also support the role of the regional ecological network in maintaining connectivity between populations.

The Dinaric Mountains, which stretch along the Adriatic Sea from Slovenia to Albania, present one of the most extensive and contiguous forests landscapes in Europe, providing crucial habitat for both pine and stone marten. In Slovenia, the Dinaric range merges with the Prealpine and Alpine foothills, forming a continuous ecological corridor that supports the distribution of both species. Pine marten is widespread in these interconnected regions, including the Dinaric, Alpine, and Prealpine zones, which is consistent with its occurrence in the neighbouring Friuli-Venezia Giulia region, Italy. On the Slovenian mainland, they tend to prefer bright, open forests, rocky habitats, and wooded riparian zones. In contrast, stone martens are more common in sub-Mediterranean areas of Slovenia—where pine martens are largely absent—and typically inhabit mixed forests [32] as well as rural and (sub)urban areas. Although ecological differences exist between the two species, they may occur sympatrically in some transitional zones [32]. In Croatia detailed distribution maps are not available, but generally it is considered both species are present in the continental and Mediterranean regions of the country, including most islands [33]. In both Slovenia and Croatia, the populations of both pine and stone marten are generally considered large and stable, so are legally regulated as game species. However, there is a lack of comprehensive scientific data on their actual population sizes, long-term trends, demographic dynamics, and fine-scale distribution, although reliable data on harvest and other mortality sources exist for Slovenia. There, in the period 2014–2024, the annual harvest of stone marten ranged from 790 individuals in 2020 to 995 in 2015, and for pine marten between 82 in 2024 and 127 individuals in 2017, respectively. During the same period, registered roadkill of stone and pine marten varied between 342+24 individuals (in 2022) and 430+49 (in 2019), respectively. In addition, between 14 and 49 stone martens and <5 pine martens were found dead each year due to other or unknown reasons [34]. No verified data on hunting bags or other causes of mortality are available in Croatia.

The primary aim of our study was to assess the genetic diversity of pine and stone martens in Croatia and Slovenia. In particular, we aimed to study the spatial distribution of genetic lineages using mitochondrial DNA control region sequences for both species Due to the limited number of reliably identified pine marten samples—mainly due to difficulties in morphological identification by non-expert collectors—we focused the analysis of population structure using microsatellite markers on stone marten, for which a sufficient number of samples was available. Nevertheless, pine martens were also included in the study to characterize mitochondrial haplotype diversity and to assess interspecific genetic differentiation.

## Materials and methods

### Samples and DNA extraction

A total of 199 stone marten and 29 pine marten samples from Croatia and Slovenia were included in this study (Fig 1 and S1 Table). Muscle samples were collected from roadkill, naturally deceased or hunted animals, by researchers, hunters and volunteers. No animal was shot or otherwise killed for the purposes of this study. Tissue samples (2 × 2 mm) were preserved in 70% ethanol, and before the DNA extraction they were air-dried under sterile conditions to remove the ethanol. The DNA was extracted using the Qiagen QIAamp® DNA kit according to the manufacturer’s instructions (Qiagen, Hilden, Germany). The DNA was eluted in the appropriate elution buffer and the sample concentration was measured with Qubit 3.0 using the Qubit dsDNA BR Assay Kit (Thermo Fisher Scientific, Waltham, MA, USA). The purity of isolated DNA was also determined using Epoch™ spectrophotometer (BioTeck, Winooski, VT, USA) measuring the 260/280 and 260/230 absorbance ratio. The extracted DNA was stored in the freezer at - 20°C until further analysis.

**Fig 1.**
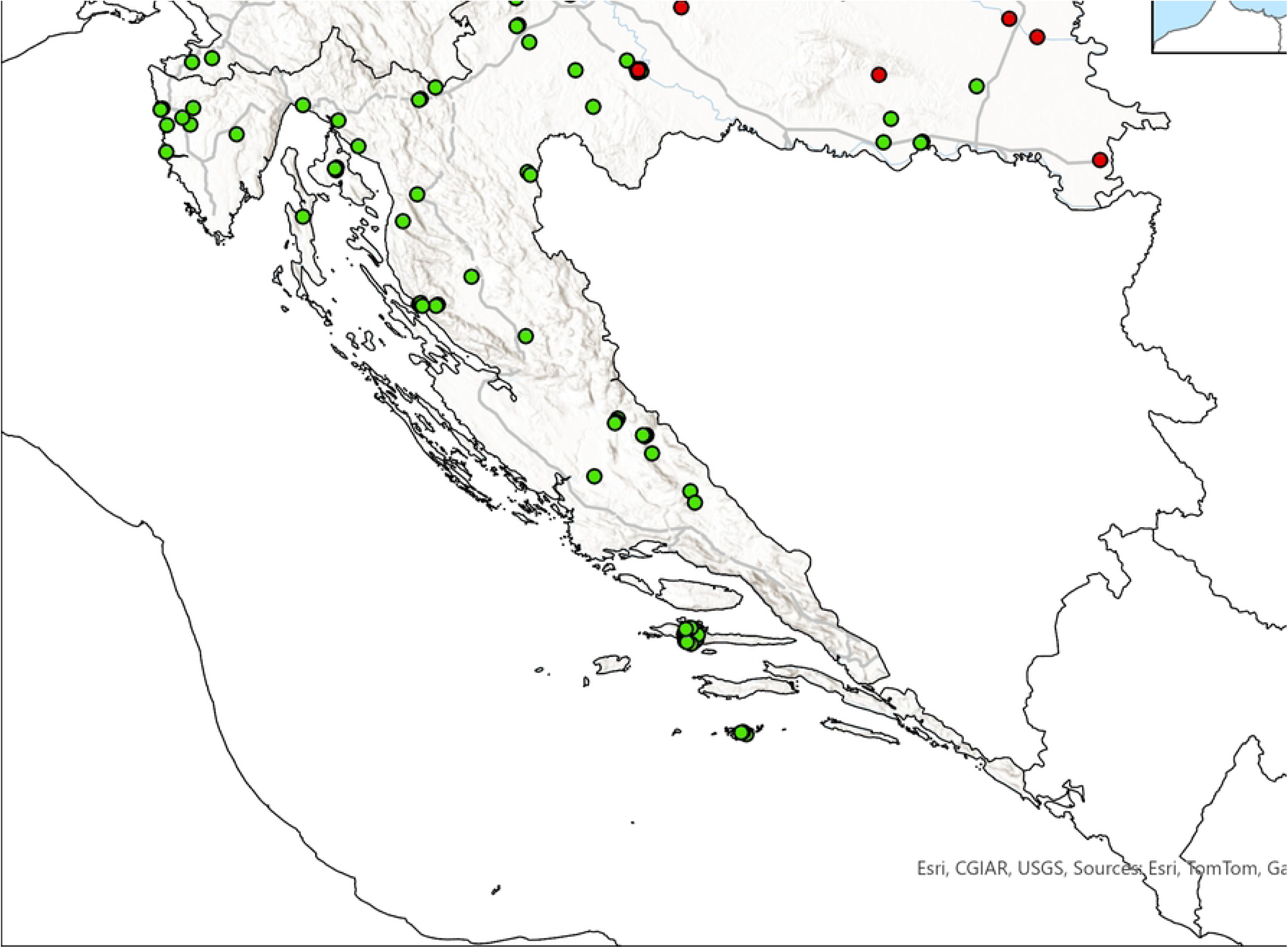
Sampling locations within the study area; detailed information on the sampled individuals and locality is provided in S1 Table.

### Mitochondrial control region amplification and sequencing

**A** partial fragment of the mitochondrial control region (CR) was amplified using the versatile primer set Dloop-MelR: 5′-ATGTCCTGTAACCATTGACTG −3′ [35] and LRCB1: 5′-TGGTCTTGTAAACCAAAAATGG −3′ [36]. All polymerase chain reactions (PCR) were performed in a 20 μl reaction mix, using DreamTaq Green PCR Master Mix (Thermo Fisher Scientific) and amplified on a Thermal Cycler 2720 (Applied Biosystems). Amplification was performed under the following conditions: incubation for 3 min at 95°C, 30 cycles with 30 s at 95°C, 45 s at 61°C and 60 s at 72°C, followed by a final extension step of 10 min at 72°C. Afterwards, 3 µL of PCR product was purified using a mix consisting of 0.2 µL of exonuclease (Exo I) (Thermo Fisher Scientific), 1 µL alkaline phosphatase (Thermo Fisher Scientific) and 2.8 µL DreamTaq Green PCR Master Mix (Thermo Fisher Scientific). The purification reaction was performed under the following conditions: 30 min at 37°C followed by 15 min at 80°C. Next, two Sanger sequencing reactions were performed in which 3 µL of purified PCR product was sequenced using a mix of 4.2 µL PCR grade water, 1.5 µL BD 5x Buffer (BD biosciences, Becton Drive Franklin Lakes, NY, USA), 1 µL of BigDye Terminator v3.1 sequencing kit (Applied Biosystems) and 0.3 µL of either Dloop-MelR or LRCB1 primer in the concentration of 10µM. The sequencing reactions were performed under the following conditions: incubation for 3 min at 96°C, 35 cycles with 10 s at 96°C, 5 s at 50°C and 4 min at 60°C. Electrophoresis of the Sanger sequencing reactions was performed on the Seq Studio Genetic Analyzer (Thermo Fisher Scientific).

### Mitochondrial genetic diversity and population structure analysis

Genetic diversity was assessed with the following parameters: (i) haplotype diversity with standard deviation (Hd ± SD), (ii) nucleotide diversity (π ± SD), (iii) number of haplotypes (h), and (iv) number of polymorphic sites (P). All parameters were estimated with the program DnaSP v.6.12 [37].

The relationship among observed and published haplotypes was evaluated by constructing median-joining haplotype networks (MJN; [38]) using PopART [39]. ArcGIS Pro (ESRI, Redlands, CA, USA) was used for geographical visualisation of data.

Since only a few samples of stone marten from Slovenia were available, the comparison between different geographical areas was conducted at the country level. This was done by calculating the pairwise fixation index (pairwise F_ST_) with 20,000 permutations to measure genetic differentiation [40,41]. Global F_ST_ and pairwise F_ST_ analyses were performed using Arlequin version 3.5.2.2 [42]. To account for different sample sizes, haplotype rarefaction curves were generated in R v4.4.1 [43] using “HaploAccum()” function in the SpideR package [44] with 1,000 permutations. Nonparametric Chao 1 estimator of total haplotype diversity was calculated using the “chaoHaplo()” function.

### Microsatellite amplification and fragment analysis

The microsatellites were amplified with the ready-to-use KAPA2G Fast Multiplex Mix (Kapa Biosystems, Wilmington, MA, USA), in accordance with the manufacturer’s instructions, using 3 µL of template DNA and 0.23 μM of final concentration for each primer used in the set. Amplification was performed under the following conditions: incubation of 3 min at 95°C, 30 cycles with 30 s at 95°C, annealing for 30 s at 55°C, extension for 1 min at 72°C followed by a final extension step of 10 min at 72°C. Fragment analysis was performed on a SeqStudio sequencer (Thermo Fischer scientific) using the GeneScan LIZ500 (−250) standard (Applied Biosystems). The results were validated with GeneMapper v.5.0 software (Applied Biosystems). We amplified 13 microsatellite loci in 3 multiplex sets containing 5, 4 and 4 microsatellites, respectively (S4 Table).

### Analysis of inter-population genetic structure and spatial patterns

The presence of null alleles and heterozygosity deficiency was assessed with FreeNA Version 1.0 [45], which uses the Expectation Maximization (EM) algorithm of Dempster et al. (1977) that accounts for deviations from Hardy-Weinberg equilibrium (HWE). Of the 13 microsatellite loci, four were removed from the analysis; Mf 8.8, due to high null allele frequencies; Mf 1.3, Mf 4.10 and Mf 8.10, due to difficulties in PCR conditions standardization and/or inconsistent results on fragment analysis.

Global F_ST_ and pairwise population differentiation (F_ST_) were assessed using Arlequin version 3.5.2.2 [42]. To investigate genetic structuring within the dataset, discriminant analysis of principal components (DAPC) was performed using the Adegenet package [46] in R v4.4.1. DAPC is a multivariate method that reduces genetic data dimensionality via principal component analysis (PCA), followed by discriminant analysis to identify genetic clusters.

For spatial population structure, Geneland v4.9.2 [47] was used within the R environment. This Bayesian method applies MCMC simulations to estimate the number of genetic populations (K) and assigns individuals to clusters based on genetic similarity, incorporating spatial autocorrelation and isolation-by-distance effects.

Finally, isolation by distance (IBD) was tested using Mantel tests, comparing genetic and geographic distances. Euclidean geographic distances were calculated in R using Adegenet, and significance was assessed through 999 Monte Carlo permutations, evaluating correlations between Edwards genetic distances and Euclidean distances.

## Results

### Mitochondrial sequence analysis

We amplified 469 bp long mtDNA CR fragments in 107 stone marten samples and 488 bp long mtDNA CR fragments from 28 out of 29 pine marten samples (S1 Table).

### Mitochondrial DNA genetic diversity of stone marten

A total of 10 haplotypes (labelled as MF1, MF2, MF5, MF6, MF7, MF8, MF9, MF10, MF11, and MF_H25) were identified in the analysed stone marten samples (4 were identified in the Slovenian samples and all 10 in the Croatian samples (S1 Table). These were compared with previously published haplotypes from European (Germany, Bulgaria, Greece, Ukraine, Russia, and Turkey) and Asian populations (Turkmenistan, Tajikistan, Russia, Kazakhstan, China, and Pakistan) obtained from GenBank. All haplotypes identified in our samples match previously published ones; thus, no novel mtDNA haplotypes were detected in either Croatia or Slovenia.

The most frequent haplotype among the analysed samples was MF2 (25.2%), followed by MF7 (24.3%) and MF11 (19.6%) (S1 Table). The haplotypes MF1, MF6, MF8, MF9, MF10, and MF_H25 were detected exclusively in Croatia, whereas MF2, MF5, MF7, and MF11 were found in both countries.

The haplotype accumulation curves by population (S1 Fig) did not reach an asymptote suggesting that further sampling is likely to increase the haplotype diversity. The value of the haplotype accumulation curve for Croatia at seven individuals is 4.2 haplotypes, while the total number of haplotypes discovered in Slovenia is four, suggesting that the inclusion of more samples from Slovenia would possibly lead to discovery of more haplotypes. The estimated Chao 1 richness for Croatia shows that 14.5 haplotypes potentially exist for this marker. The Chao 1 richness for Slovenia was 6.0 haplotypes. The low estimate is likely due to the limited sequences (n=7).

The total number of polymorphic sites was nine. The overall haplotype diversity was 0.811 ± 0.015 and nucleotide diversity was 0.005 ± 0.001. The Croatian group showed higher nucleotide (π = 0.006 ± 0.001) and haplotype diversity (Hd = 0.814 ± 0.015) compared to the Slovenian group (π = 0.005 ± 0.002 and Hd = 0.810 ± 0.039, respectively) (Table 1) but the possible reason for this result is the limited sample size in Slovenia, which is also reflected in the lower estimate of Chao 1 richness and the lower number of total haplotypes observed. Tajima’s D test yielded a positive value for the Croatian group, which may suggest recent population expansion, and a negative value for the Slovenian group, potentially indicating genetic drift. However, as none of the result were statistically significant, definitive conclusions cannot be drawn—particularly for the Slovenian population, where the small sample size limits interpretative power.

**Table 1.** Genetic variation in the analysed samples of stone marten.

| | P | h | Hd (SD) | $\pi$ (SD) | Tajima's D |
| --- | --- | --- | --- | --- | --- |
| Overall<br>(n=107) | 9 | 10 | 0.811 (0.015) | 0.005 (0.001) | 1.528 (p>0.10) |
| Slovenia<br>(n=7) | 6 | 4 | 0.810 (0.039) | 0.005 (0.002) | -0.339 (p>0.10) |
| Croatia<br>(n=100) | 9 | 10 | 0.814 (0.015) | 0.006 (0.001) | 1.540 (p>0.10) |

P: number of polymorphic sites, h: number of haplotypes, Hd: haplotype diversity, $\pi$ nucleotide diversity, SD: standard deviation.

### Insight into phylogeographic structure of stone marten from mitochondrial haplotype network

A total of 85 stone marten mtDNA CR haplotypes (truncated to the overlapping 469 bp) were included in the median-joining network (S1 and S2 Tables), suggesting two major haplogroups with a clear geographical division (Fig 2). One haplogroup consists primarily of our samples along with those from eastern Turkey, while the other, known as the Major Eurasian group, includes our samples together with those from Western, Eastern, and Northern Europe, Western Turkey, and Asia. Despite the well-documented geographical division between Western and Eastern Turkey, as previously described by Arslan et al. and Ishii et al., no clear geographical structuring is observed within the Major Eurasian group [24,27]. The cluster containing haplotypes from Eastern Turkey also includes four novel haplotypes from Croatia (MF6, MF8, MF9, and MF10), along with the common haplotypes MF_H1 and MF_H15, which is shared by samples from Eastern Turkey and Slovenia. Within the Major Eurasian group, the majority of Croatian samples belong to the MF-H26 haplotype, while the remaining samples are distributed among MF-1, MF_Hap6, and MF_Hap9. A smaller number of samples belong to haplotype MF-Hap6, which is also shared with samples from Greece and Bulgaria. Samples from Slovenia were assigned to haplotypes MF-1, MF-Hap9, and MF_H26. Interestingly, MF-1 and MF-Hap9 also include samples from Russia and Ukraine—despite these regions are geographically distant from the current study area. The MF_H26 haplotype has a wider distribution, occurring in southern Germany, Bulgaria, Greece, and western Turkey. The star-like topology in the Major Eurasian group is observed around the common haplotype MF_H26, with most haplotypes closely related by a single mutation step. The intermediate haplotype MF-1 also exhibits a slightly weaker star-like structure, with four mutation steps connecting it to the Eastern Turkey cluster and three mutational steps from the central haplotypes in the second cluster MF_H26.

**Fig 2.**
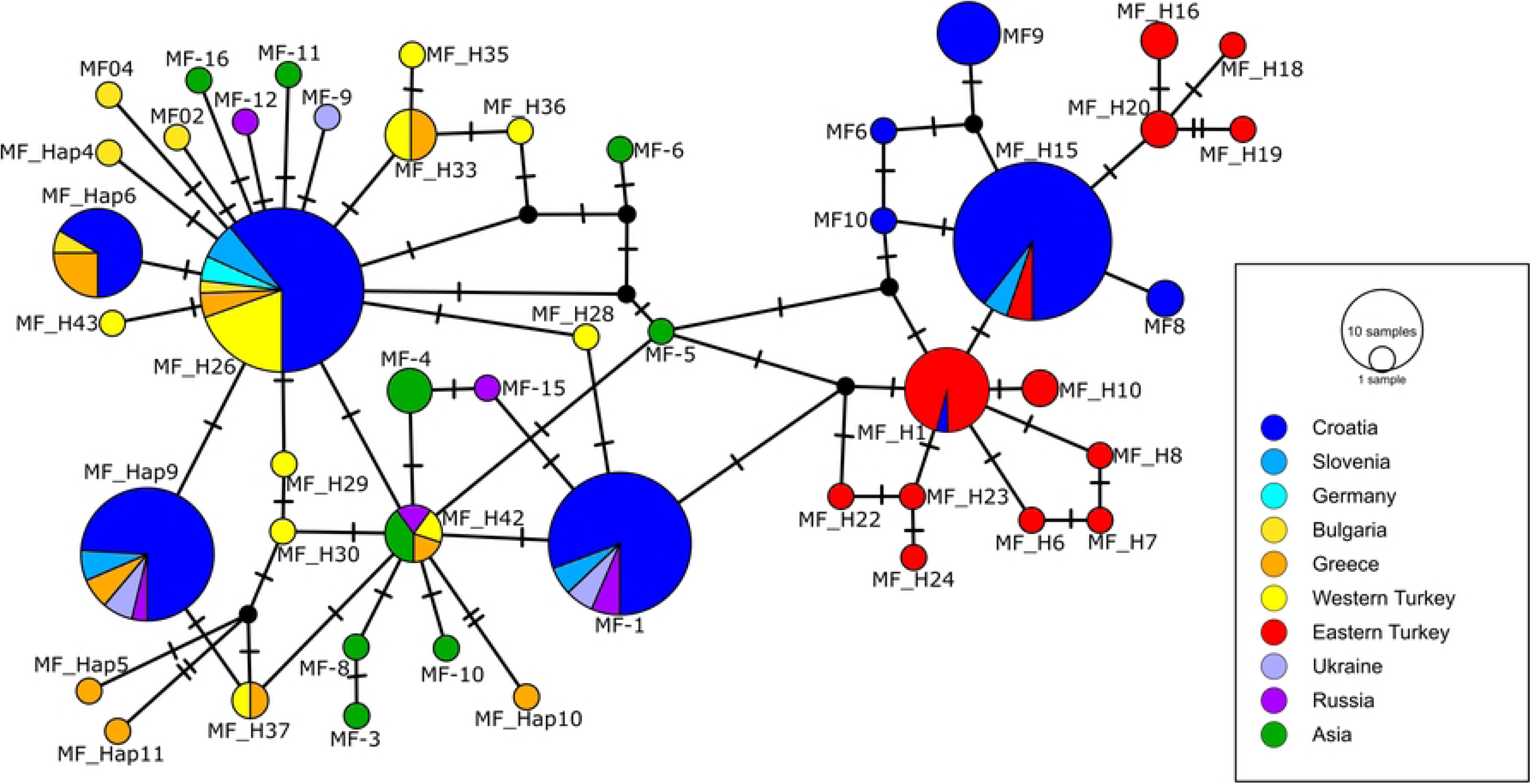
Median-joining haplotype network for 85 mtDNA control region sequences (469 bp) of stone marten (S1 and S2 Tables). Each circle indicates a haplotype (the name is given next to the circle), with the size of the circle proportional to the number of individuals bearing that haplotype. Slash marks in lines connecting the circles indicate the number of nucleotide substitutions between adjacent haplotypes. Black dots indicate intermediate haplotypes not detected. The colour of each circle indicates the country or geographical region (in the case of Turkey) where each haplotype was found. We used the same colour for these regions as Ishii et al. [24].

The overall haplotype diversity (Hd) was high (0.919 ± 0.008), with a relatively low nucleotide diversity (π) of 0.007 ± 0.001. The Major Eurasian and Eastern Turkey groups showed slightly lower haplotype diversities of 0.860 ± 0.015 and 0.864 ± 0.019, respectively, with identical nucleotide diversities (0.004 ± 0.001) among each other (Table 2). Tajima’s D values varied between groups, being negative overall (−1.123, p>0.10) and in the Eastern Turkey group (−0.980, p>0.10), and positive in the Major Eurasian group (1.791, p>0.05). In contrast to the results for Tajima’s D, which were all not statistically significant, Fu’s Fs values were strongly negative across all groups, with highly significant values observed overall (−32.055, p<0.001), for the Major Eurasian group (−22.643, p<0.001), and for the Eastern Turkey group (−6.786, p=0.001).

**Table 2.**
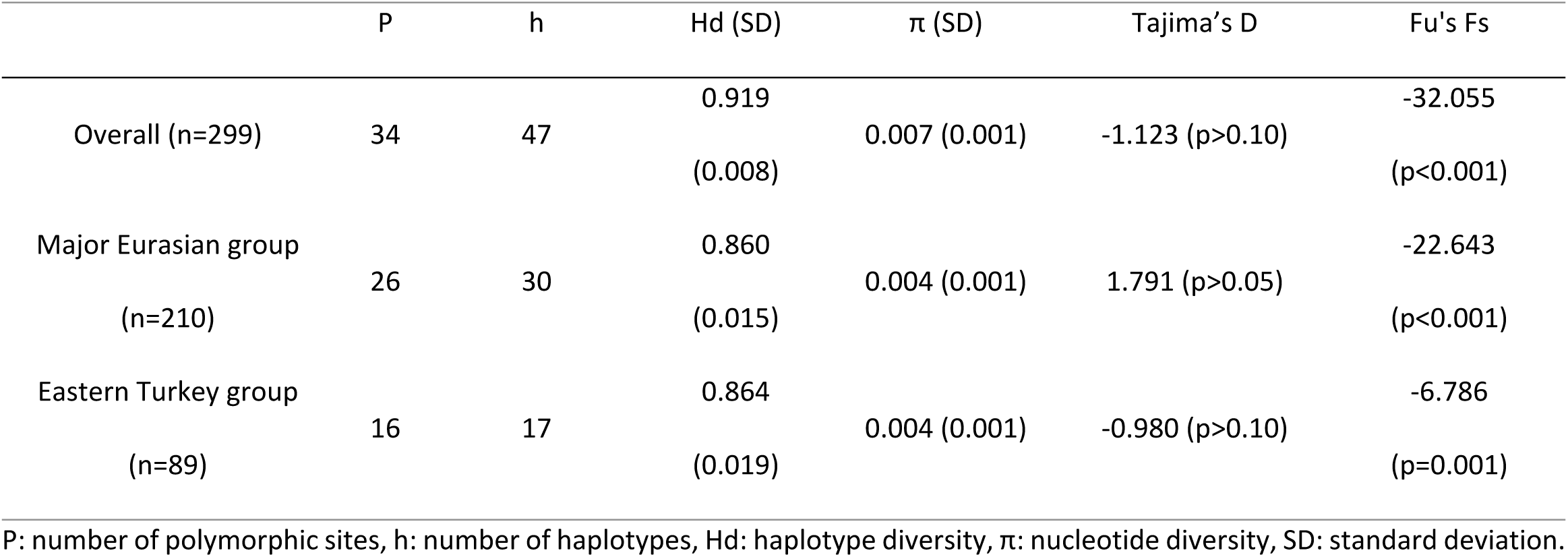
Molecular diversity indices based on mitochondrial control region sequences for stone marten.

| | P | h | Hd (SD) | $\pi$ (SD) | Tajima's D | Fu's Fs |
| --- | --- | --- | --- | --- | --- | --- |
| Overall (n=299) | 34 | 47 | 0.919<br>(0.008) | 0.007 (0.001) | -1.123 (p>0.10) | -32.055<br>(p<0.001) |
| Major Eurasian group<br>(n=210) | 26 | 30 | 0.860<br>(0.015) | 0.004 (0.001) | 1.791 (p>0.05) | -22.643<br>(p<0.001) |
| Eastern Turkey group<br>(n=89) | 16 | 17 | 0.864<br>(0.019) | 0.004 (0.001) | -0.980 (p>0.10) | -6.786<br>(p=0.001) |

P: number of polymorphic sites, h: number of haplotypes, Hd: haplotype diversity, $\pi$ : nucleotide diversity, SD: standard deviation.

### Spatial genetic structure of stone marten

The DAPC plot based on individual multilocus genotypes revealed clear genetic structuring of the stone marten populations (Fig 3), with individuals divided into 6 clusters. Samples from the island of Hvar (HR Hvar) formed a well-defined cluster, distinctly separated along the first discriminant axis from the majority of mainland individuals, indicating strong genetic divergence. A separation of individuals from the islands of Krk and Lastovo from Hvar was also evident, although they overlapped with the mainland populations, suggesting moderate gene flow or shared ancestry. Samples from Slovenia and the Croatian mainland showed substantial genetic admixture and formed a central, more diffuse cluster. These results support the presence of restricted gene flow and possible founder effects or genetic drift acting in the Hvar population. The number and strength of discriminant functions are visualized in the DA eigenvalue plot (inset), confirming the first axis as the most informative in explaining between-group variance.

**Fig 3.**
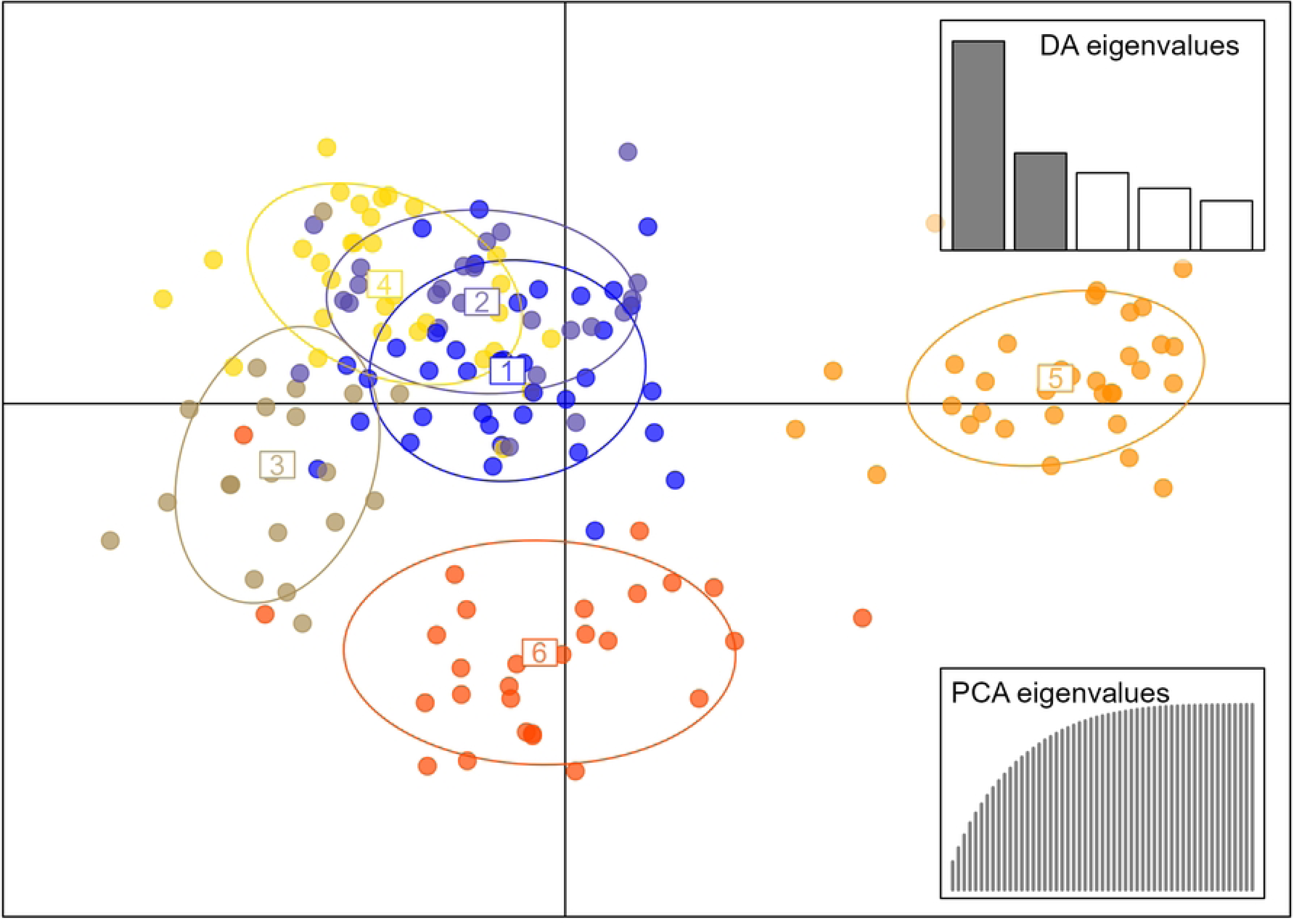
Analysis of population structure of stone marten in Croatia and Slovenia using DAPC. The main graph plots show the first two discriminant axes (explaining 15.1% and 7.6% of the variation, respectively). Clusters are shown by different colours and shapes, while points represent individuals. Bar plot shows assignment of individuals to the six genetic clusters without prior information.

The spatial genetic structure inferred from the Geneland analysis identified four distinct genetic clusters, each showing strong geographical association (Fig 4). These spatial patterns are broadly consistent with the results of the DAPC analysis, where island populations (particularly Hvar) formed clearly separated genetic groups, while mainland populations exhibited more admixture. Clusters 1 and 2 of the Geneland output largely correspond to individuals from mainland regions, particularly those that were assigned to admixed or overlapping clusters in the DAPC plot. Cluster 3, whose high posterior probabilities are centred on a southern region, likely reflects a geographically isolated population—possibly corresponding to individuals on or near the southern Adriatic islands (e.g., Lastovo), where unique genetic signatures were detected in the DAPC analysis. Cluster 4, which spans northern and central areas, also includes individuals that formed intermediate groupings in the DAPC space.

**Fig 4.**
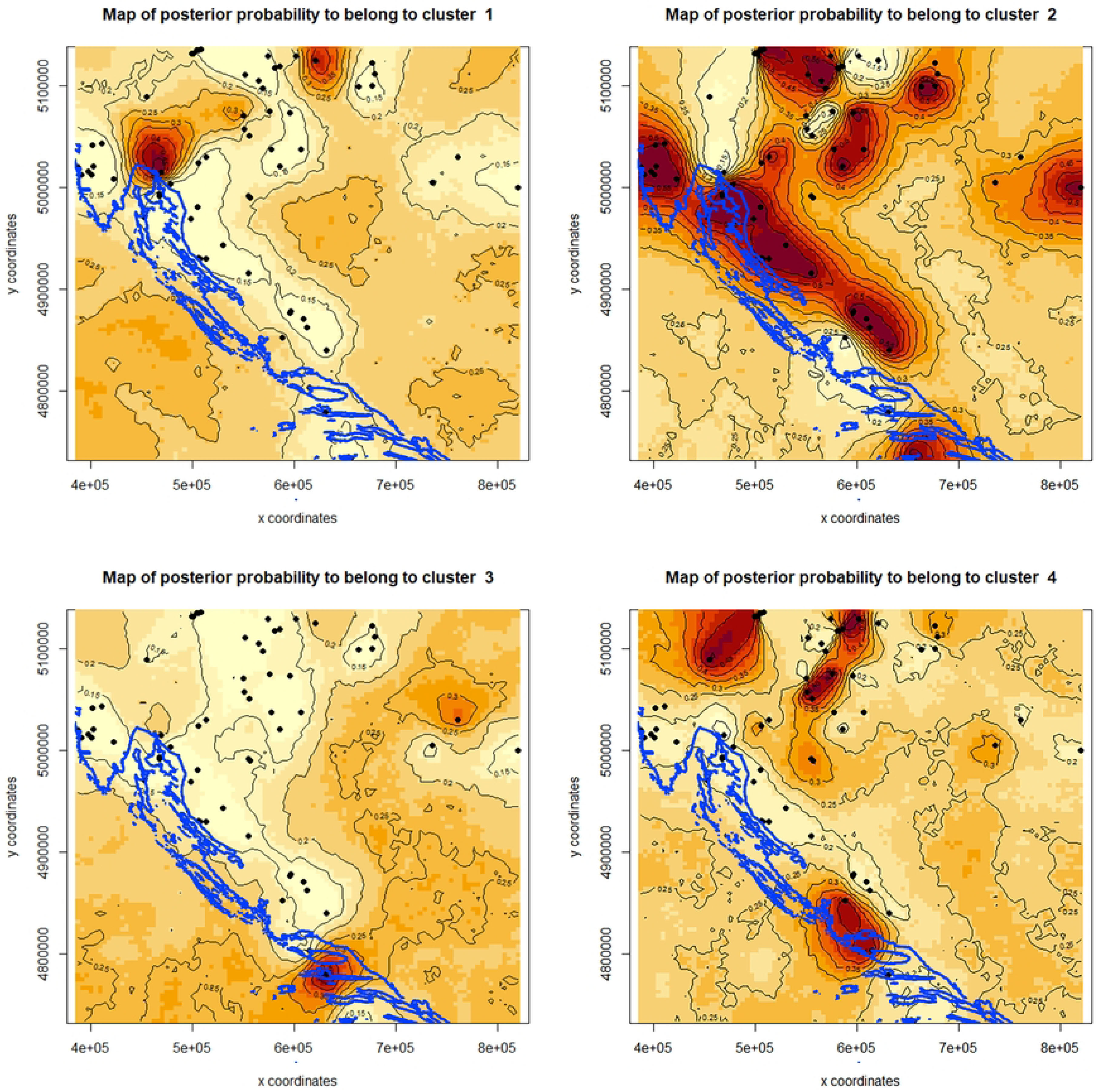
Map showing spatially explicit maps of posterior probabilities for assignment to each of the four inferred genetic clusters, based on Geneland analysis using an independent allele frequencies model that accounts for potential null alleles. The analysis included samples from Croatia and Slovenia (represented by black points), with darker areas indicating higher probabilities of cluster membership. The Adriatic coast is marked by a blue line.

The results of the pairwise F_ST_ analysis (Table 3) show a very high genetic differentiation between the samples from the island of Hvar and the rest of the samples, with the highest differentiation with the samples from the island of Lastovo (0.546, p<0.05). The samples from the island of Lastovo also showed high genetic differentiation with the samples from the mainland.

**Table 3.** Pairwise F_ST_ between clusters/populations, as defined in Fig 3.

|  | Hvar | Krk | Lastovo | Mainland HR |
| --- | --- | --- | --- | --- |
| Krk | **0.442** |  |  |  |
| Lastovo | **0.546** | 0.145 |  |  |
| Mainland HR | **0.264** | 0.0150 | **0.188** |  |
| Mainland SI | **0.312** | 0.0354 | **0.219** | 0.005 |

\* Values in bold indicate significant differences with $\alpha = 0.05$

Taken together, the pairwise F_ST_, Geneland and DAPC analyses reveal a consistent pattern of geographic population structure, with island populations showing strong genetic differentiation and mainland populations forming broader, less distinct clusters. These findings suggest limited gene flow between island and mainland populations and historical isolation of stone martens inhabiting remote islands.

### Isolation by distance

Microsatellite-based genetic distances (Fig 5) were not correlated with geographical distances among populations (observation = 0.235, p = 0.133).

**Fig 5.**
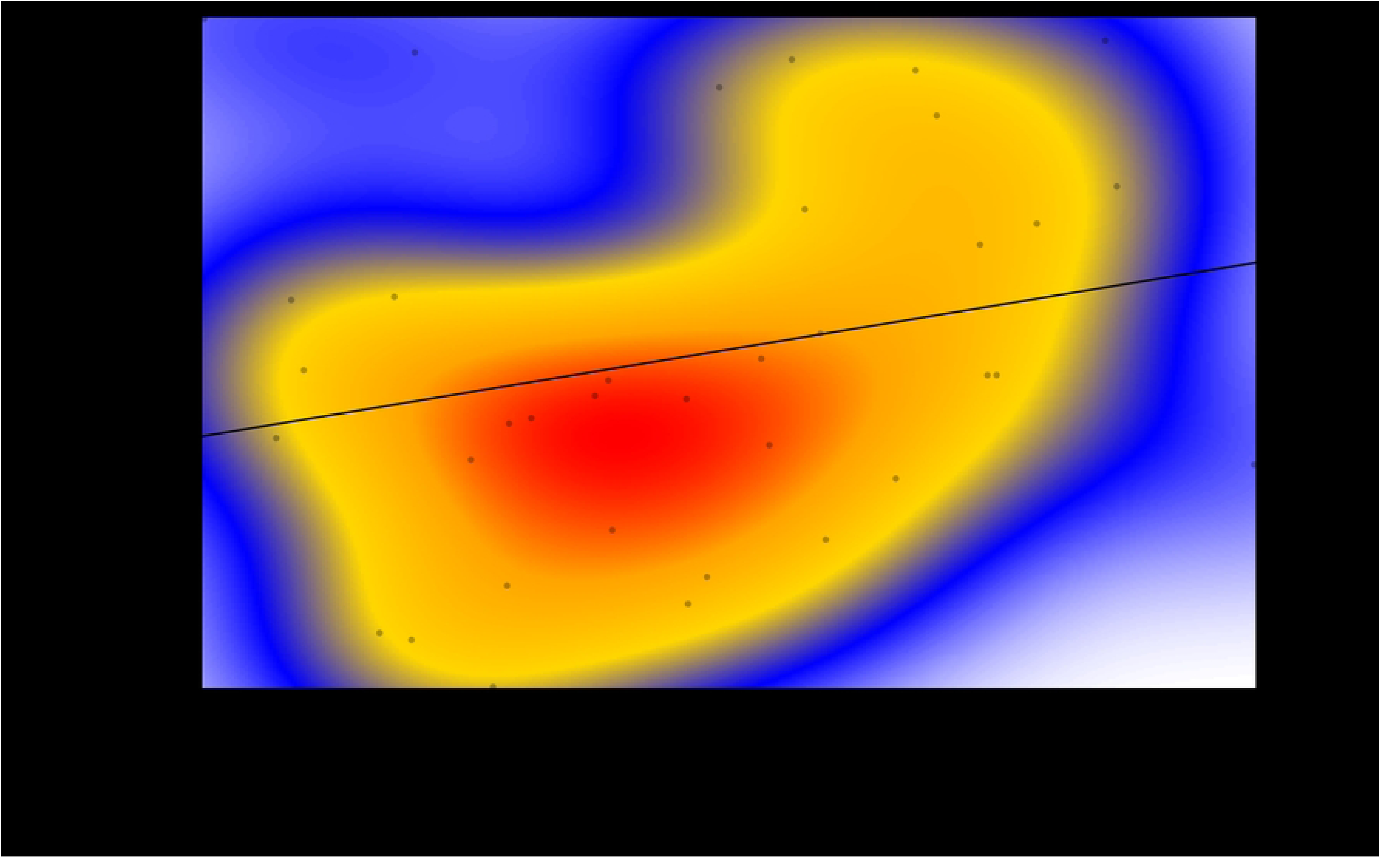
Plot obtained for the isolation by distance analysis. Local density of points was plotted using a two-dimensional kernel density estimation. Dgeo: geographical distances in degrees; Dgen: genetic distance.

### Mitochondrial DNA genetic diversity of pine marten

The alignment of partial mitochondrial control region sequences (488 bp) from 28 pine martens—one from Slovenia and 27 from Croatia—revealed eight haplotypes. Four of these haplotypes (MM19, MM5, MM43, and MM44) had been previously described by Ruiz-González et al., while the remaining four (MM2, MM3, MM5, and MM6) were identified as novel in this study [29]. Due to the limited sample size— comprising only one individual from Slovenia and 27 from Croatia—we assessed genetic diversity across the combined sample set rather than by country. The analysis revealed a total of nine variable sites, representing 1.8% of the total 488 bp mitochondrial control region sequence. Overall haplotype diversity was 0.828 ± 0.042, and nucleotide diversity was 0.007 ± 0.001, indicating moderate mitochondrial genetic diversity within the sampled individuals.

### Insight into phylogeographic structure of pine marten from mitochondrial haplotype network

A median-joining haplotype network was constructed using a total of 161 haplotypes (Fig 6), including both newly generated sequences and those retrieved from GenBank (S1 and S2 Tables). Haplotypes previously deposited in GenBank were named according to their original designations (S2 Table). The network analysis supports the phylogenetic structure described by Ruiz-González et al., identifying two major haplogroups within *M. martes*: the Fennoscandian–Russian (FNR) group and a broader cluster comprising nearly the entire European range of the species [29]. This latter group is further subdivided into two phylogeographic lineages: the Mediterranean (MED) and the Central–Northern European (CNE) groups. Our samples were assigned to both groups; however, all novel haplotypes identified in this study fall into the CNE lineage.

**Fig 6.**
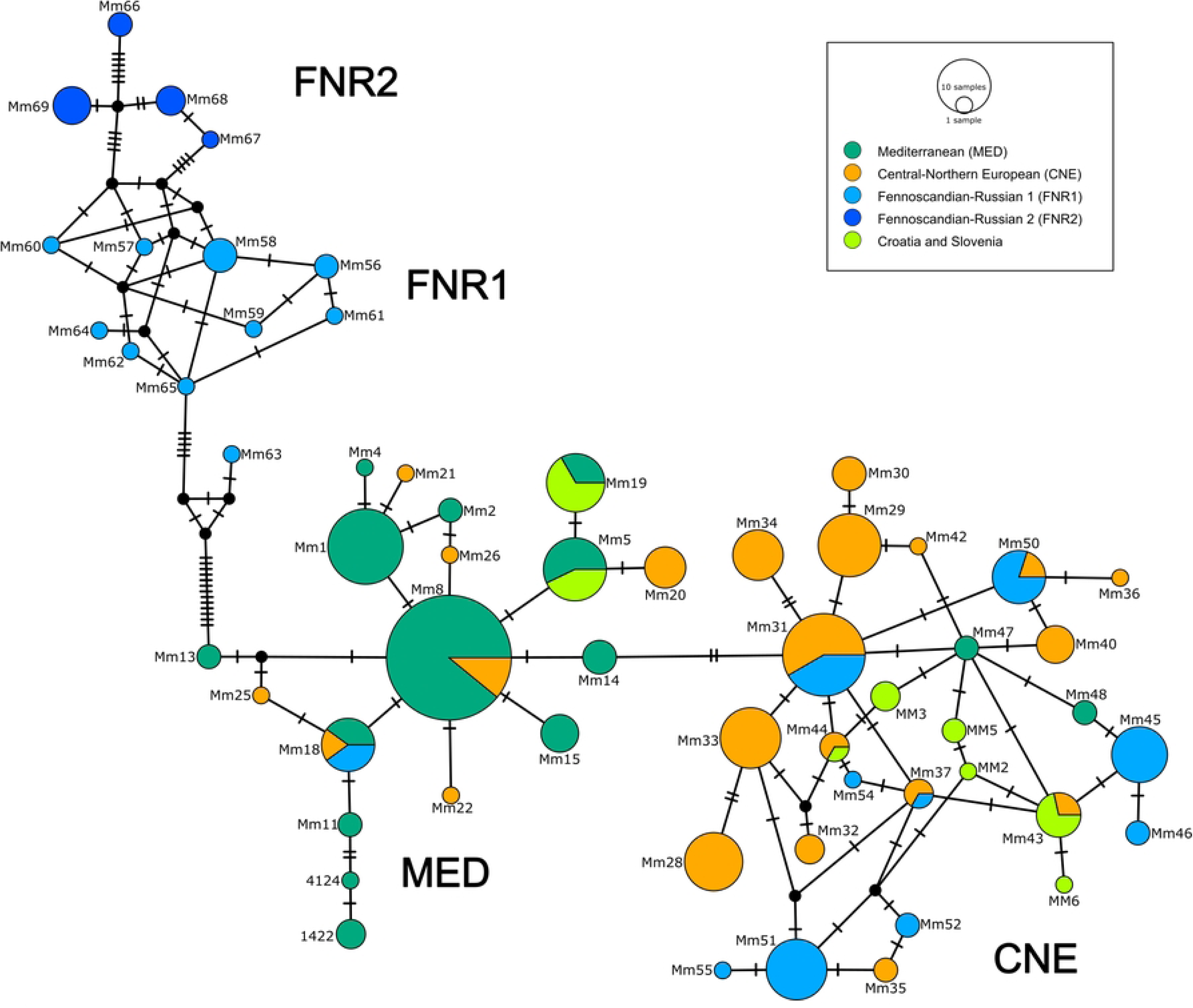
The median-joining network of mtDNA CR haplotypes from the European pine marten populations is colour-coded according to phylogroups. The number of mutations (greater than one) between haplotypes is represented by perpendicular lines. Circle size is proportional to the frequency of each haplotype. For haplotype designations and distributions, see S2 Table.

Molecular diversity indices were calculated using the haplotype sequences from this study and previously published data (Table 4 and S2 Table). Overall haplotype diversity (Hd) was high at 0.935 ± 0.008, with nucleotide diversity (π) of 0.0135 ± 0.001. Among the three phylogeographic groups, the Central– Northern European group showed high haplotype diversity (0.932 ± 0.008) but relatively low nucleotide diversity (0.007 ± 0.001). The Fennoscandian–Russian group had similarly high haplotype diversity (0.930 ± 0.030) with a slightly higher nucleotide diversity (0.012 ± 0.001). In contrast, the Mediterranean group exhibited the lowest values for haplotype and nucleotide diversity, at 0.770 ± 0.031 and 0.003 ± 0.001, respectively.

**Table 4.** Molecular diversity indices based on mitochondrial control region sequences for pine marten.

| | P | h | Hd (SD) | $\pi$ (SD) | Tajima's D | Fu's $F_s$ |
| --- | --- | --- | --- | --- | --- | --- |
| Overall |  |  | 0.935 | 0.0135 | -0.224 | -19.370 |
| (n=318) | 45 | 55 | (0.008) | (0.001) | ($p>0.10$) | ($p=0.003$) |
| CNE |  |  | 0.932 | 0.007 | -0.240 | -9.992 |
| (n=152) | 20 | 27 | (0.008) | (0.001) | ($p>0.10$) | ($p=0.007$) |
| FNR |  |  | 0.930 | 0.012 | 0.568 | -2.664 |
| (n=25dn) | 19 | 14 | (0.030) | (0.001) | ($p>0.10$) | ($p=0.132$) |
| MED |  |  | 0.770 | 0.003 | -1.350 | -9.939 |
| (n=141) | 16 | 18 | (0.031) | (0.001) | ($p>0.10$) | ($p=0.001$) |

P: number of polymorphic sites, h: number of haplotypes, Hd: haplotype diversity, $\pi$ : nucleotide diversity, SD: standard deviation.

Tajima’s D values were negative in all groups except the FNR group, which showed a positive but non-significant value (0.568, p>0.10). Although none of the Tajima’s D values reached the threshold of statistical significance, the consistently negative values in the overall dataset and in the CNE and MED groups may indicate potential population expansion. This interpretation is further supported by the significantly negative Fu’s Fs values observed in the overall dataset (−19.370) as well as in CNE (−9.992) and MED (−9.939) groups. The FNR group, however, showed a smaller and non-significant Fu’s Fs value (−2.664). These findings are consistent with patterns previously reported by Ruiz-González et al., who also identified signals of demographic expansion and historical structuring among *M. martes* mitochondrial lineages across Europe [29]. Regardless of the statistically non-significant results for Tajima’s D, the observation of a combination of high haplotype diversity and relatively low nucleotide diversity and the highly negative and significant values of Fu’s Fs (Fu 1997) support the hypothesis of a potential recent expansion in the stone marten, which is particularly visible in the Major Eurasian haplogroup [48]. Support for this hypothesis was also reported by Arslan et al. and Ishii et al. [24,27].

## Discussion

This study provides new insights into the genetic diversity and phylogeographic structure of two sympatric mustelid species, pine marten and stone marten, in a biogeographically complex region spanning Slovenia and Croatia. By integrating mitochondrial and microsatellite data, we evaluated spatial genetic patterns and demographic processes reflecting both natural and anthropogenic influences.

### Mitochondrial diversity and phylogeography

The mitochondrial haplotype network identified two well-supported stone marten phylogenetic groups: one aligning with the widely distributed Major Eurasian lineage, and the other that clusters with haplotypes from eastern Turkey. This division is congruent with the mitochondrial phylogeny previously reported by Arslan et al., who described two distinct clades: one restricted to eastern Turkey and another encompassing western Turkey, Greece, and Bulgaria [27]. Haplotypes from Croatia (MF6, MF8, MF9, and MF10) were clustered with the eastern group, suggesting a deep genetic divergence in the Balkan area, likely reflecting historical isolation in glacial refugia. The grouping of these haplotypes with eastern Anatolian lineages is consistent with findings by Ishii et al., who also proposed the existence of multiple glacial refugia in Anatolia and the Balkans that contributed to the current diversity and distribution of stone marten in Europe [24]. The presence of widespread haplotypes such as MF_H26 and MF-1 in Slovenia and Croatia indicates postglacial expansion and gene flow from the Balkans across the continent [27]. This pattern also highlights Turkey as an important refugium for the stone marten during the Last Glacial Maximum [22]. Our results further support the idea that the refugial population on the Balkan Peninsula expanded into western Turkey, large parts of Europe, and Central Asia after the LGM, facilitated by the absence of significant geographical barriers. In addition, there is evidence for to the existence of two distinct refugia in Turkey during the LGM, separated by the Taurus Mountains [24]. It is assumed that populations from the western refugium migrated to central and western Europe, but also to south-eastern European countries (e.g., Greece, Bulgaria), in the postglacial period.

Such patterns of multiple refugial origins, complex phylogeographic structures, and range expansions have also been documented in other mustelids, including European badger (*Meles meles*) [49], polecat (*Mustela putorius*) [50], least weasel (*Mustela nivalis*) [51,52], and pine marten [29]. All studies reinforce the idea that south-eastern Europe serves as a complex contact zone where divergent lineages of several mustelid species can coexist and potentially hybridise.

The pine marten haplotypes from Croatia and Slovenia correspond to the main haplogroups defined by Ruiz-González et al. and confirm the presence of two main mitochondrial lineages: the Mediterranean and the Central–Northern European clades [5]. All novel haplotypes detected in our study fall within the CNE group, supporting their broad postglacial expansion and suggesting previously undocumented genetic variation in the Alps and the Dinaric region. Following the topology of the phylogenetic tree of Ruiz-González et al. the clustering of the pine marten samples reflects common ancestry with populations from Central and Eastern Europe, rather than the more differentiated Mediterranean clade [29].

### Demographic trends and historical processes

The negative Fu’s Fs values detected for both stone and pine marten (particularly in the CNE clade for pine marten and in the Major Eurasian group for stone marten) suggest population expansion after the Last Glacial Maximum. Although Tajima’s D values were not significant, the combination of mtDNA diversity and network structure supports scenarios of demographic growth, as previously proposed in broader phylogeographic studies. These expansion signals are especially relevant for understanding the role of the Balkans and Anatolia as refugia and source areas for the recolonization of both marten species across Europe.

### Population structure and spatial differentiation of stone marten

Microsatellite data revealed a moderate genetic structuring in the stone marten, with distinct clusters corresponding to island populations (e.g. Hvar) and more admixed mainland groups. The DAPC analysis showed a strong separation of insular populations along the first discriminant axis, while individuals from the Croatian and Slovenian mainland formed overlapping clusters, suggesting continuous gene flow and limited barriers to dispersal. This finding is consistent with other studies showing that anthropogenic landscapes, rather than topographic complexity, shape genetic structuring in mesocarnivores [20,53,54]. The Geneland spatial analysis further confirmed the presence of four genetic clusters with geographical coherence. While one cluster was clearly associated with the southern Adriatic region (island of Hvar), others showed a broader distribution across central and northern areas. This pattern indicates both localized genetic drift in isolated populations and broad-scale gene flow between connected regions. Remarkably, the Dinaric Mountains, traditionally considered a geographical barrier, have not disrupted the genetic connectivity of stone martens, but may even have facilitated their movement, as has been observed in other mesocarnivores in alpine and subalpine habitats. These differentiation patterns are consistent with the findings of Wereszczuk et al., who reported a moderate population structure in stone marten across central-eastern Europe [21]. Their study revealed significant F_ST_ values between regions in Poland and eastern Germany, likely reflecting historical fragmentation and current habitat discontinuities. Although our F_ST_ values are similarly moderate, the geographical structuring differs. While Wereszczuk et al. identified a broader east–west division and reduced connectivity within agricultural landscapes, our data revealed fine-scale differentiation between islands and mainland sites, emphasising the natural isolation of island populations and relatively unrestricted gene flow across the rugged mainland terrain [21]. Collectively, these results emphasize the flexibility of stone marten in navigating heterogeneous landscapes, even though local population structures may evolve through geographical or ecological isolation. In addition, the Mantel test showed that isolation by distance was not statistically significant (p = 0.15), suggesting that geographical distance alone cannot explain the observed genetic differentiation in the populations studied. This further supports the influence of landscape features, such as insularity or habitat discontinuities, over purely spatial separation. The absence of IBD may result from two overlapping processes: widespread gene flow across mainland Croatia and Slovenia, and the presence of a fine-scale population structure. Both processes deviate from expectations of IBD and together likely explain its absence in this system.

Although our results provide valuable insights into the genetic diversity and structure of martens in the northern Dinaric and Adriatic regions, several limitations must be acknowledged. The most notable is the limited sample size for pine martens, particularly in Slovenia, which restricts the power of inferences regarding phylogeographic structure and demographic history. Furthermore, the use of only a partial mitochondrial marker and a relatively small set of microsatellite loci, while informative, does not capture the full genomic complexity of these species. Future research should therefore apply genome-wide approaches, such as whole-genome resequencing, to obtain a higher-resolution picture of population connectivity, local adaptation, and evolutionary history. Such genomic tools will be especially important for resolving fine-scale structure, detecting subtle admixture, and understanding the functional basis of adaptation in response to both natural and anthropogenic pressures. Integrating ecological data with genomic analyses will also enhance our ability to inform conservation and management strategies for these sympatric carnivores in a changing landscape.

## Conclusions

This study provides new insights into the genetic structure and phylogeographic history of stone and pine martens in south-eastern Europe, a region of recognized biogeographic complexity. By analysing mitochondrial DNA and microsatellites, we identified significant genetic diversity and distinct lineage patterns shaped by postglacial expansion and historical isolation. The presence of both widespread and regionally restricted haplotypes underscores the role of the Balkans (and Anatolia, as revealed earlier by other authors) as important glacial refugia. Microsatellite data revealed a moderate population structure of the stone marten, particularly highlighting the genetic distinctiveness of island populations in Croatia. However, overall connectivity between mainland populations remains high, and the lack of significant isolation by distance suggests that geographical barriers such as the Dinaric Mountains do not substantially limit gene flow in this species.

Despite these new findings, our study also highlights the need for more comprehensive genomic approaches to better resolve population history, admixture, and adaptive variation. Future studies should incorporate genome-wide markers and integrate environmental data to deepen our understanding of the evolutionary and ecological processes shaping marten populations in this heterogeneous region.

## Author contributions

**Conceptualization:** Elena Buzan

**Formal analysis:** Luka Duniš, Tilen Komel

**Funding acquisition:** Elena Buzan, Boštjan Pokorny, Magda Sindičić

**Methodology:** Elena Buzan, Luka Duniš, Tilen Komel

**Project administration:** Elena Buzan

**Resources:** Boštjan Pokorny, Magda Sindičić, Zoran Marčić

**Supervision:** Elena Buzan

**Validation:** Elena Buzan, Luka Duniš, Tilen Komel

**Visualization:** Luka Duniš

**Writing – Original Draft Preparation:** Elena Buzan

**Writing – Review & Editing:** Elena Buzan, Boštjan Pokorny, Magda Sindičić, Carlos Rodríguez Fernandes, Tilen Komel, Luka Duniš

## Acknowledgments

We thank all the collaborators and founders who help accelerate collaboration with hunters as citizen scientists in wildlife monitoring and biodiversity research. We thank many hunters, foresters and other colleagues for collecting and providing the samples.

## Data Accessibility

All sequence data submitted to NCBI’s GenBank database is also automatically mirrored in the European Nucleotide Archive (ENA) and the DNA Data Bank of Japan (DDBJ).

## Supporting information

**S1 Fig. Haplotype accumulation curves.** Haplotype accumulation curves by population of stone marten (*Martes foina*) samples from the present study from Croatia (red line) and Slovenia (cyan line).

**S1 Table. Sample data.** Basic data on pine marten (*Martes martes*) and stone marten (*Martes foina*) included in the study. The “Haplotype” column represents the recognized haplotype, the “Haplotype – network” column represents how the haplotypes are labelled in the haplotype networks (Figs 2 and 6).

**S2 Table. *Martes foina* haplotypes obtained from GenBank.** List of *Martes foina* haplotypes retrieved from GenBank. The “Haplotype - GenBank” column represents the haplotype names obtained from GenBank, the “Haplotype – network” column represents how the haplotypes are labelled in the haplotype network (Fig 2).

**S3 Table. *Martes martes* haplotypes obtained from GenBank.** List of *Martes foina* haplotypes retrieved from GenBank. The “Haplotype - GenBank” column represents the haplotype names obtained from GenBank, the “Haplotype – network” column represents how the haplotypes are labelled in the haplotype network (Fig 2).

**S4 Table. Microsatellite loci primers.** Microsatellite loci analyzed in fragmentation analysis of pine marten (*M. Martes*) and stone marten (*M. foina*) published by Basto et al. 2010.

## References

1. Proulx G, Aubry K, Birks J, Buskirk S, Fortin C, Frost H, et al. World distribution and status of the genus *Martes* in 2000. In: Harrison DJ, Fuller AK, Proulx G, editors. Martens and fishers (Martes) in human-altered environments. New York: Springer-Verlag; 2005. pp. 21–76. doi:10.1007/0-387-22691-5_2

2. Larroque J, Ruette S, Vandel J, Devillard S. Where to sleep in a rural landscape? A comparative study of resting sites pattern in two syntopic *Martes* species. Ecography. 2015;38: 1129–1140. doi:10.1111/ecog.01133

3. IUCN. Martes foina: Abramov, A.V., Kranz, A., Herrero, J., Choudhury, A. & Maran, T.: The IUCN Red List of Threatened Species 2016: e.T29672A45202514. 2015. doi:10.2305/IUCN.UK.2016-1.RLTS.T29672A45202514.en

4. Mitchell-Jones TJ. The atlas of European mammals. London: T. & A. D. Poyser Academic press, on behalf of the Societas Europaea mammalogica; 1999.

5. Ruiz-González A, Madeira MJ, Randi E, Urra F, Gómez-Moliner BJ. Non-invasive genetic sampling of sympatric marten species (Martes martes and Martes foina): assessing species and individual identification success rates on faecal DNA genotyping. Eur J Wildl Res. 2013;59: 371–386. doi:10.1007/s10344-012-0683-6

6. Tikhonov, A., Cavallini, P., Maran, T., Krantz, A., Herrero, J., Giannatos, G., et al. Vormela peregusna. In: IUCN 2012. IUCN Red List of Threatened Species. Version 2012.2. 2008. Available: www.iucnredlist.org

7. Herr J, Schley L, Roper TJ. Stone martens (*Martes foina*) and cars: Investigation of a common human-wildlife conflict. European Journal of Wildlife Research. 2009;55: 471–477. doi:10.1007/s10344-009-0263-6

8. Virgos E., Zalewski A., Rosalino L., Mergey M. Habitat ecology of genus *Martes* in Europe: a review of the evidences. Biology and Conservation of Martens, Sables, and Fishers. Cornell University Press, New York; 2012. pp. 255–266.

9. Balestrieri A, Ruiz-González A, Capelli E, Vergara M, Prigioni C, Saino N. Pine marten vs. stone marten in agricultural lowlands: a landscape-scale, genetic survey. Mamm Res. 2016;61: 327–335. doi:10.1007/s13364-016-0295-8

10. Balestrieri A, Remonti L, Ruiz-González A, Gómez-Moliner BJ, Vergara M, Prigioni C. Range expansion of the pine marten (*Martes martes*) in an agricultural landscape matrix (NW Italy). Mammalian Biology. 2010;75: 412–419. doi:10.1016/j.mambio.2009.05.003

11. Mergey M, Helder R, Roeder J-J. Effect of forest fragmentation on space-use patterns in the European pine marten (*Martes martes*). Journal of Mammalogy. 2011;92: 328–335. doi:10.1644/09-MAMM-A-366.1

12. Vergara M, Cushman SA, Ruiz-González A. Ecological differences and limiting factors in different regional contexts: landscape genetics of the stone marten in the Iberian Peninsula. Landscape Ecol. 2017;32: 1269–1283. doi:10.1007/s10980-017-0512-0

13. Herrero, J., Kranz, A., Skumatov, D., Abramov, A.V., Maran, T., Monakhov, V.G. Martes martes: The IUCN Red List of Threatened Species 2016: e.T12848A45199169. 2015. doi:10.2305/IUCN.UK.2016-1.RLTS.T12848A45199169.en

14. Domingo-Roura X. Genetic distinction of marten species by fixation of a microsatellite region. Journal of Mammalogy. 2002;83: 907–912. doi:10.1644/1545-1542(2002)083<0907:GDOMSB>2.0.CO;2

15. Vercillo F, Livia L, Nadia M, Bernardino R, Ettore R, Fausto P. A simple and rapid PCR-RFLP method to distinguishing *Martes martes* and *Martes foina*. Conservation Genetics. 2004;5: 869–871. doi:10.1007/s10592-004-1866-9

16. Colli L, Cannas R, Deiana AM, Gandolfi G, Tagliavini J. Identification of mustelids (Carnivora: Mustelidae) by mitochondrial DNA markers. Mammalian Biology. 2005;70: 384–389. doi:10.1016/j.mambio.2005.02.005

17. Pilot M, Gralak B, Goszczyński J, Posłuszny M. A method of genetic identification of pine marten (Martes *martes*) and stone marten (*Martes foina*) and its application to faecal samples. Journal of Zoology. 2007;271: 140–147. doi:10.1111/j.1469-7998.2006.00179.x

18. Ruiz-González A, Rubines J, Berdión O, Gómez-Moliner BJ. A non-invasive genetic method to identify the sympatric mustelids pine marten (*Martes martes*) and stone marten (*Martes foina*): preliminary distribution survey on the northern Iberian Peninsula. Eur J Wildl Res. 2008;54: 253–261. doi:10.1007/s10344-007-0138-7

19. Croose E, Birks JDS, O’Reilly C, Turner P, Martin J, MacLeod ET. Sample diversity adds value to non-invasive genetic assessment of a pine marten (*Martes martes*) population in Galloway Forest, southwest Scotland. Mamm Res. 2016;61: 131–139. doi:10.1007/s13364-015-0257-6

20. Ruiz-González A, Gurrutxaga M, Cushman SA, Madeira MJ, Randi E, Gómez-Moliner BJ. Landscape genetics for the empirical assessment of resistance surfaces: the European pine marten (*Martes martes*) as a target-species of a regional ecological network. Maldonado JE, editor. PLoS ONE. 2014;9: e110552. doi:10.1371/journal.pone.0110552

21. Wereszczuk A, Leblois R, Zalewski A. Genetic diversity and structure related to expansion history and habitat isolation: stone marten populating rural–urban habitats. BMC Ecol. 2017;17: 46. doi:10.1186/s12898-017-0156-6

22. İbiş O, Koepfli K, Özcan S, Tez C. Genetic analysis of Turkish martens: Do two species of the genus *Martes* occur in Anatolia? Zoologica Scripta. 2018;47: 390–403. doi:10.1111/zsc.12289

23. Tsoupas A, Andreadou M, Papakosta MA, Karaiskou N, Bakaloudis DE, Chatzinikos E, et al. Phylogeography of *Martes foina* in Greece. Mammalian Biology. 2019;95: 59–68. doi:10.1016/j.mambio.2019.02.004

24. Ishii H, Amaike Y, Nishita Y, Abramov AV, Masuda R. Phylogeography of the stone marten (*Martes foina*: Mustelidae: Mammalia) in Eurasia, based on a mitochondrial DNA analysis. Mamm Res. 2023;68: 375–381. doi:10.1007/s13364-023-00690-6

25. Nagai T, Raichev EG, Tsunoda H, Kaneko Y, Masuda R. Preliminary study on microsatellite and mitochondrial dna variation of the stone marten *Martes foina* in Bulgaria. Mammal Study. 2012;37: 353–358. doi:10.3106/041.037.0410

26. Vergara M, Basto MP, Madeira MJ, Gómez-Moliner BJ, Santos-Reis M, Fernandes C, et al. Inferring population genetic structure in widely and continuously distributed carnivores: The stone marten (*Martes foina*) as a case study. PLoS ONE. 2015;10. doi:10.1371/journal.pone.0134257

27. Arslan Y, Demiϊrtaş S, Herman JS, Pustilnik JD, Searle JB, Gündüz İ. The Anatolian glacial refugium and human-mediated colonization: a phylogeographical study of the stone marten (*Martes foina*) in Turkey. Biological Journal of the Linnean Society. 2020;129: 470–491. doi:10.1093/biolinnean/blz180

28. Nishita Y, Väinölä R, Abramov AV, Masuda R. Diversity of MHC class II *DRB* alleles and mitochondrial DNA in northern and eastern European pine marten, *Martes martes* (Mammalia: Mustelidae). Biological Journal of the Linnean Society. 2023; blad133. doi:10.1093/biolinnean/blad133

29. Ruiz-González A, Madeira MJ, Randi E, Abramov AV, Davoli F, Gómez-Moliner BJ. Phylogeography of the forest-dwelling European pine marten (M*artes martes*): new insights into cryptic northern glacial refugia: European pine marten phylogeography. Biol J Linn Soc Lond. 2013;109: 1–18. doi:10.1111/bij.12046

30. Basto MP, Santos-Reis M, Simões L, Grilo C, Cardoso L, Cortes H, et al. Assessing genetic structure in common but ecologically distinct carnivores: the stone marten and red fox. Somers C, editor. PLoS ONE. 2016;11: e0145165. doi:10.1371/journal.pone.0145165

31. Kyle CJ, Davison A, Strobeck C. Genetic structure of European pine martens (Martes martes), and evidence for introgression with M. americana in England. 2003.

32. Kryštufek B. Sesalci Slovenije. Ljubljana: Prirodoslovni muzej Slovenije; 1991.

33. Janicki, Z., Slavica, A., Konjević, A., Severin, K. Zoologija divljači. Zavod za patologiju i uzgoj divljači. Sveučilište u Zagrebu, Veterinarski fakultet; 2007.

34. OSLIS - Central Slovenian Hunting Information System. Slovenian Forestry Institute; 2016 [cited 2025 Jun 30]. Available from: http://oslis.gozdis.si/

35. Sato JJ, Yasuda SP, Hosoda T. Genetic Diversity of the Japanese Marten (*Martes melampus*) and Its Implications for the Conservation Unit. Zoological Science. 2009;26: 457–466. doi:10.2108/zsj.26.457

36. Davison A, Birks JDS, Brookes RC, Messenger JE, Griffiths HI. Mitochondrial phylogeography and population history of pine martens *Martes martes* compared with polecats *Mustela putorius*. Molecular Ecology. 2001;10: 2479–2488. doi:10.1046/j.1365-294X.2001.01381.x

37. Librado P, Rozas J. DnaSP v6.11: A software for comprehensive analysis of DNA polymorphism data. Bioinformatics. 2017;25: 1451–1452. doi:10.1093/bioinformatics/btp187

38. Bandelt HJ, Forster P, Rohl A. Median-joining networks for inferring intraspecific phylogenies. Molecular Biology and Evolution. 1999;16: 37–48. doi:10.1093/oxfordjournals.molbev.a026036

39. Leigh JW, Bryant D. POPART : full-feature software for haplotype network construction. Nakagawa S, editor. Methods Ecol Evol. 2015;6: 1110–1116. doi:10.1111/2041-210X.12410

40. Reynolds J, Weir BS, Cockerham CC. Estimation of the coancestry coefficient: Basis for a short-term genetic distance. Genetics. 1983;105: 767–779.

41. Slatkin M. A measure of population subdivision based on microsatellite allele frequencies. Genetics. 1995;139: 457–462.

42. Excoffier L, Lischer HEL. Arlequin suite ver 3.5: A new series of programs to perform population genetics analyses under Linux and Windows. Molecular Ecology Resources. 2010;10: 564–567. doi:10.1111/j.1755-0998.2010.02847.x

43. R Core Team (2025). R: a language and environment for statistical computing. R foundation for statistical computing, Vienna, Austria. 2025.

44. Brown SDJ, Collins RA, Boyer S, Lefort M, Malumbres-Olarte J, Vink CJ, et al. S PIDER : An R package for the analysis of species identity and evolution, with particular reference to DNA barcoding. Molecular Ecology Resources. 2012;12: 562–565. doi:10.1111/j.1755-0998.2011.03108.x

45. Chapuis MP, Estoup A. Microsatellite null alleles and estimation of population differentiation. Molecular Biology and Evolution. 2007;24: 621–631. doi:10.1093/molbev/msl191

46. Jombart T. Adegenet: A R package for the multivariate analysis of genetic markers. Bioinformatics. 2008;24: 1403–1405. doi:10.1093/bioinformatics/btn129

47. Guillot G, Estoup A, Mortier F, Cosson JF. A spatial statistical model for landscape genetics. Genetics. 2005;170: 1261–1280. doi:10.1534/genetics.104.033803

48. Grant WS, Bowen BW. Shallow population histories in deep evolutionary lineages of marine fishes: Insights from sardines and anchovies and lessons for conservation. Journal of Heredity. 1998. pp. 415–426. doi:10.1093/jhered/89.5.415

49. Frantz AC, Pope LC, Carpenter PJ, Roper TJ, Wilson GJ, Delahay RJ, et al. Reliable microsatellite genotyping of the Eurasian badger (*Meles meles*) using faecal DNA. Molecular Ecology. 2003;12: 1649–1661. doi:10.1046/j.1365-294X.2003.01848.x

50. Watanabe R, Nishita Y, Abramov AV, Peeva S, Raichev E, Masuda R. Phylogeography of *Mustela eversmanii* and *M. putorius* (Carnivora: Mustelidae) throughout continental Eurasia, based on mitochondrial control-region sequences. Mamm Res. 2025 [cited 17 Jul 2025]. doi:10.1007/s13364-025-00795-0

51. Kurose N, Abramov AV, Masuda R. Comparative phylogeography between the ermine *Mustela erminea* and the least weasel *M. nivalis* of palaearctic and nearctic regions, based on analysis of mitochondrial dna control region sequences. Zoological Science. 2005;22: 1069–1078. doi:10.2108/zsj.22.1069

52. Sato T, Abramov AV, Raichev EG, Kosintsev PA, Väinölä R, Murakami T, et al. Phylogeography and population history of the least weasel (*Mustela nivalis*) in the Palearctic based on multilocus analysis. J Zool Syst Evol Res. 2020;58: 408–426. doi:10.1111/jzs.12330

53. Torre I, Pulido T, Vilella M, Díaz M. Mesocarnivore distribution along gradients of anthropogenic disturbance in mediterranean landscapes. Diversity. 2022;14: 133. doi:10.3390/d14020133

54. Soccorsi AE, LaPoint SD. Assessing spatiotemporal patterns of mesocarnivores along an urban-to-rural gradient. J Wildl Manag. 2023;87: e22474. doi:10.1002/jwmg.22474

